# The retinal light response is modulated by an mGluR5-mediated retrograde signal from ON-bipolar cells to photoreceptors

**DOI:** 10.1101/2025.11.13.687902

**Authors:** Catherine W. Morgans, Tammie L. Haley, Gaoying Ren, Ian A. M. Taylor, Kate L. Randall, Robert M. Duvoisin

## Abstract

The ON visual pathway is initiated by the deactivation of mGluR6, coupled to the opening of TRPM1 channels in retinal ON-bipolar cell dendrites. Here, we show that a second metabotropic glutamate receptor, mGluR5, is localized with TRPM1 and mGluR6 in the dendrites of ON-bipolar cells. To examine the function of mGluR5, we performed electroretinogram (ERG) recordings and found that the amplitude of the b-wave, which is primarily a measure of ON-bipolar cell light-driven activity, is reduced in mGluR5 knock-out mice compared to wild type. In the mGluR5^-/-^ retina, we observed weaker mGluR6 immunofluorescence in the dendritic tips of ON-bipolar cells that could explain the smaller ERG b-wave. To observe the effect of mGluR5 without perturbing mGluR6 expression, wild type mice were injected with MTEP, an allosteric antagonist of mGluR5. MTEP increased the amplitude of the b-wave in response to dim stimuli and caused an inflection in the intensity-response plot for flashes in the mesopic range. In the brain, postsynaptic mGluR5 regulates presynaptic glutamate release via endocanabinoid-mediated retrograde signaling. Therefore, we tested the effect of the CB1 receptor antagonist, SR1417A, on the ERG and found that the b-wave was affected as by MTEP, including an inflection in the intensity-response. We further showed that the CB1 receptor agonist, ACEA, reversed the effects of MTEP. Together, our results indicate that mGluR5 plays a role in gain-control at the photoreceptor to ON-bipolar cell synapses, likely via an endocannabinoid-mediated retrograde feedback.

**Significance Statement:** Light-ON stimuli are transmitted from photoreceptors to ON-bipolar cells via three pathways, which are used under different light intensities, from dim to bright light. For each pathway, glutamate is the neurotransmitter released by rods and cones, and the postsynaptic role of the mGluR6-TRPM1 signaling cascade in ON-bipolar cells is well studied, with defects known to cause congenital stationary night blindness. We show here that mGluR5 is also present in ON-bipolar cell dendrites and regulates synaptic transmission between photoreceptors and ON-bipolar cells. In particular, we find that mGluR5 facilitates the transition between the different photoreceptor to ON-bipolar cell pathways via mGluR5-controlled retrograde release of endocannabinoids.

## Introduction

Parallel ON and OFF visual pathways, which signal increases and decreases in light intensity, are established at the first synapse in the visual system between photoreceptors and bipolar cells and are preserved throughout all subsequent levels of visual processing. Vertebrate photoreceptors are depolarized in the dark, which supports the tonic release of glutamate from their synaptic terminals; light stimulation causes photoreceptors to hyperpolarize, thereby reducing glutamate release (Marc and Lam, 1981; Bloomfield and Dowling, 1985; Copenhagen and Jahr, 1989). The ON and OFF responses of bipolar cells are determined by the type of glutamate receptors they express. The OFF-bipolar cell response is mediated by ionotropic receptors, which cause the cell to hyperpolarize in response to the light-induced decrease in synaptic glutamate (DeVries, 2000; DeVries et al., 2006; Borghuis et al., 2014; Lindstrom et al., 2014; Ichinose and Hellmer, 2016). In contrast, the ON-bipolar cell response is initiated by a unique G protein-coupled receptor (GPCR), mGluR6 (Nomura et al., 1994; Vardi et al., 2000), which causes a reversal of response polarity at this synapse, such that deactivation of mGluR6 following a drop in synaptic glutamate results in the opening of TRPM1 cation channels, to generate a depolarizing response to light (Morgans et al., 2009; Shen et al., 2009; Koike et al., 2010). Typical of signaling via GPCRs, the ON-bipolar cell signal transduction pathway is built on a cascade of protein-protein interactions with the potential for modulation at every stage (Martemyanov and Sampath, 2017). Cone photoreceptors connect to both ON- and OFF-bipolar cells in the outer plexiform layer (OPL) of the retina; in contrast, in mammals, rod photoreceptors connect to rod bipolar cells, which generate ON-type responses that are fed into the retinal ON-circuitry and inhibit the OFF-circuitry via AII amacrine cells in the inner plexiform layer (IPL).

In addition to mGluR6, several groups have shown that a group I mGluR (either mGluR1 or mGluR5) is also present in ON-bipolar cells (Koulen et al., 1997 (rat); Tagawa et al., 1999 (mouse); Sen and Gleason, 2006 (chicken); Joselevitch et al., 2007 (goldfish); Klooster et al., 2009 (zebrafish); Rampino and Nawy, 2011 (mouse); Hozumi et al., 2013 (rat); Hellmer et al., 2018 (mouse)). Here, we show that mGluR5, but not mGluR1, is found in mouse ON-bipolar cell dendrites. Electroretinogram (ERG) recordings of wild type and mGluR5^-/-^ mice reveal that mGluR5 signaling modulates the ERG b-wave, suggesting that this receptor regulates synaptic gain between photoreceptors and bipolar cells.

In the brain, mGluR5 modulates synaptic transmission through retrograde endocannabinoid signaling (Kano et al., 2009). mGluR5 activates phospholipase C, generating diacylglycerol (DAG), which can be hydrolyzed by diacylglycerol lipase (DAGL), leading to the formation of 2-arachidonoyl glycerol (2-AG). 2-AG can diffuse across the synapse to activate presynaptic cannabinoid (CB1 or CB2) receptors and inhibit neurotransmitter release (Varma et al., 2001; Araque et al., 2017; Augustin and Lovinger, 2018). Interestingly, CB1 receptors (CB1Rs) have recently been shown to regulate glutamate release from rod bipolar cells in the inner retina (Estay et al., 2024). Here, we explored whether such a mechanism occurs at photoreceptor synapses and found a strong effect of CB1 activity on the ERG-b-wave consistent with mGluR5-driven release of endocannabinoids from ON-bipolar cells at scotopic and mesopic intensities.

## Materials and Methods

### Animals and Chemicals

This study was carried out in accordance with the principles of the Basel Declaration, and the animal use protocol was approved by the Institutional Animal Care and Use Committee at Oregon Health & Science University. C57BL/6 wild type (WT), as well as mGluR1 and mGluR5 knockout (KO) mice (Conquet et al., 1994; Lu et al., 1997) were used for this study. All mice had access to food (PicoLab Rodent Diet 20, no. 5053; PMI Nutrition International, St. Louis, MO, USA) and water *ad libitum* throughout the study. Housing facilities were maintained at 20 – 21°C and kept on a 12 h light:12 h dark cycle. Mice were monitored daily to ensure good general health throughout the study.

Chemicals were purchased from Fisher Scientific (Pittsburgh, PA). The mGluR5 allosteric antagonist, 3-((2-Methyl-4-thiazolyl)ethynyl)pyridine (MTEP), was obtained from Hello Bio (Princeton, NJ). The CB1 agonist arachidonyl-2’-chloroethylamide (ACEA) was purchased from Cayman Chemical (Ann Arbor, MI) and the CB1 antagonist/inverse agonist, SR14176A, from Tocris Bioscience (Bio-Techne, Minneapolis, MN) and resuspended in Cremophor (Selleck Chemicals, Houston, TX): ethanol : PBS (1:1:18).

### RNAscope and Immunohistochemistry

The anterior portion of the eye and lens were removed, and the posterior eyecups were fixed in 4% (w/v) paraformaldehyde in phosphate-buffered saline (PBS; pH 7.4) for 30 – 60 min (immunohistochemistry) or 24 hrs (RNAscope, Bio-Techne) at 4°C. The eyecups were cryoprotected by immersion in sucrose-PBS solutions of increasing osmolality (2 hrs, 10%; 2 hrs, 20%; and overnight in 30%), then embedded in OCT (Sakura Finetek, Torrance, CA). 20 μm transverse cryostat sections were thaw-mounted onto Superfrost/Plus slides (Fisher Scientific) and stored frozen.

### Immunohistochemistry

After thawing to room temperature, eyecup sections were permeabilized and blocked by incubation at room temperature for 1 hr in antibody incubation solution (AIS: 3% (v⁄v) normal horse serum, 0.5% (v⁄v) Triton X-100, 0.025% (w⁄v) NaN_3_ in PBS). Sections were then incubated for 1 hr at room temperature with primary antibodies diluted in AIS. After washing with PBS three times, sections were incubated in the secondary antibodies (goat anti-mouse conjugated to Alexa Fluor 488 and goat anti-rabbit-Alexa Fluor 594; Jackson ImmunoResearch, West Grove, PA) diluted in AIS for 1 hr at room temperature. Slides were washed again in PBS and cover-slipped with Crystal⁄Mount mounting medium (Biomeda, Foster City, CA). Primary antibodies were obtained as follows: monoclonal mouse anti-mGluR1 from BD Bioscience (San Jose, CA; catalog #556389), rabbit anti-mGluR5 antiserum from Spring Biosciences (Pleasanton, CA; catalog #E2870), rabbit monoclonal anti-mGluR5 from Cell Signaling Technologies (Danvers, MA; catalog #55920), rabbit anti-protein kinase Cα (PKCα) from MilliporeSigma (Burlington, MA; catalog # P4334). Two rabbit monoclonal anti-CB1R antibodies were used, one from Cell Signaling Technologies (CST antibody; catalog # 93815) and one from Synaptic Systems (SySy antibody; Goettingen, Germany; catalog # 258 008) both of which have been validated on CB1R-knockout tissue (Li et al., 2020; Steiger et al., 2023). The mouse monoclonal anti-mGluR6 antibody was a generous gift from Melina Agosto (Dalhousie University, Halifax, Canada) and has been previously validated (Agosto et al., 2021). Antibodies to RGS7 and RGS11 were kind gifts of Theodore Wensel and have been described previously (Morgans et al., 2007). The sheep anti-TRPM1 antibody was kindly provided by Kirill Martemyanov (The Scripps Research Institute, Jupiter, FL) and previously validated (Cao et al., 2011). Peanut agglutinin (PNA) conjugated to Alexa-488 was from Molecular Probes (Eugene, OR; catalog # L-21409).

Fluorescence images were acquired with a Leica TCS SP8 X confocal microscope using a X60/1.42 oil immersion objective. Image brightness and contrast were enhanced using Image J (imagej.net).

To examine mGluR5 and mGluR6 co-localization, high magnification images of mGluR5 and mGluR6 double labeling were deconvolved using Leica Lightning software. Pearson’s colocalization coefficients were calculated using the EzColocalization plugin (Stauffer et al., 2018) in ImageJ to compare the distributions of mGluR5 and mGluR6 immunofluorescence in the outer plexiform layer after thresholding with Costes’s method. For rod bipolar cells, the average Pearson’s coefficient was calculated from single optical sections through 14 regions of the OPL containing only rod spherules. For cone-bipolar cells, the average Pearson’s coefficient was calculated from single optical sections from 18 individual cone pedicles.

To quantify differences in the immunofluorescence for mGluR6, TRPM1, RGS7, and RGS11 between WT and mGluR5-KO mouse retina, regions of interest (ROIs) were obtained from WT and KO paired retina sections using ImageJ. Then, the corrected total cell fluorescence (CTCF) was calculated based on the integrated signal density, as well as the background intensity randomly selected outside the signals (CTCF = Integrated Signal Density – (ROI Area x Background Fluorescence)). The areas of ROIs in both genotypes are the same. The ratio of CTCF_KO_ /CTCF_WT_ was calculated. A ratio near 1 indicates no change in protein expression in KO with respect to WT, whereas a ratio below 1 indicates reduced expression in the KO. To minimize the impact of bleaching on our results, image acquisition parameters were optimized during live continuous imaging on an area adjacent to the ROI, followed by imaging the ROI, and paired KO and WT retina sections were imaged with the same gain and laser intensity and the same number of optical sections. Statistical significance was calculated with a ratio t-test. Data are reported as mean ± SEM.

### RNAscope

RNAscope in situ hybridization was performed according to the manufacturer’s protocol (Bio-Techne) with the following exceptions and alterations: no antigen retrieval step was performed; protease III was used at an incubation time of 18 minutes at room temperature; all wash steps were performed on the slide within wells made by an ImmEdge pen (Vector Laboratories, Newark, CA).

### Electroretinogram recordings

Mice were dark-adapted overnight and prepared for recording under dim red light between Zeitgeber time (ZT) 5 and ZT7. Anesthesia was initiated via i.p. injection of ketamine:xylazine (100:10 mg/kg) and maintained with booster injections (30:3 mg/kg) at approximately 30 min intervals until completion of the experiment (generally 40 – 60 minutes). Body temperature was maintained between 36.5 – 38°C by placing the mouse on a water-circulated heating pad monitored with a rectal thermometer. Before recording, the pupils were dilated with 2.5% phenylephrine and 1% tropicamide, and the cornea was anesthetized with 1% topical proparacaine. A wire loop placed under the upper teeth was used to draw the mouse into a custom-made holder which stabilized the head and allowed delivery of 95% O_2_ /5% CO_2_ (carbogen; ∼0.25 l/min) to minimize breathing artifacts during recording. The recording was made using a platinum wire placed against the cornea with a small drop of 2.5% hypromellose to prevent drying. A loop electrode placed around the eye served as a reference electrode and a needle electrode placed in the tail served as a ground. The mice were placed in a Ganzfeld sphere, and light stimuli were provided by a custom-designed 502 nm LED panel. Light stimulus intensity was controlled with neutral density filters and by altering flash duration, and was measured post-hoc with an ILT5000 radiometer (International Light, Newburyport, MA) with a scotopic filter. Traces were recorded with customized software (ERGlab, Richard Weleber, Casey Eye Institute, Portland, OR).

Full-field scotopic electroretinograms (ERGs) from both eyes were simultaneously recorded after a series of flashes ranging from -3.93 to 2.41 log(cd•s/m^2^). For dim intensities (-3.93 to -2.5 log(cd•s/m^2^)), ERG recordings to 8-12 flashes were averaged. For mid-range intensities (-2.5 to 0 log(cd•s/m^2^)), three trials were averaged with inter-flash intervals of 20 seconds. For bright intensities (0 to 2.41 log(cd•s/m^2^)), two trials were averaged with inter-flash intervals of 30 to 120 seconds. Photopic ERGs were recorded on a steady 74 cd/m^2^ white light background.

ERGlab data was exported for offline processing using a custom Python script and the SciPy package (https://docs.scipy.org/doc). The start of the flash stimulus was set to time zero and trace baselines were adjusted such that the 100 ms prior to the flash were zero. Control and experimental data sets were compared using a Student t-test in SigmaPlot 15 software (Systat, San Jose, CA). Data are reported as mean ± SEM.

## Results

### mGluR5 is localized to bipolar cell dendrites

There are multiple reports of group I, Gq-coupled mGluRs expressed in bipolar cells in the outer plexiform layer (OPL; (Koulen et al., 1997; Tagawa et al., 1999; Sen and Gleason, 2006; Joselevitch et al., 2007; Klooster et al., 2009; Rampino and Nawy, 2011; Hozumi et al., 2013; Hellmer et al., 2018). As a first step to determine if mGluR1 or mGluR5 transcripts are present in bipolar cells, we queried single-cell transcriptome databases. Mouse retina single-cell RNA sequencing studies (Shekhar et al., 2016; Li et al., 2024) reveal co-expression of mGluR6 and mGluR5 in most ON-bipolar cell types, as well as mGluR5 transcripts in OFF BC1B cells, that don’t express mGluR6 (Fig. 1*A*).

**Figure 1.**
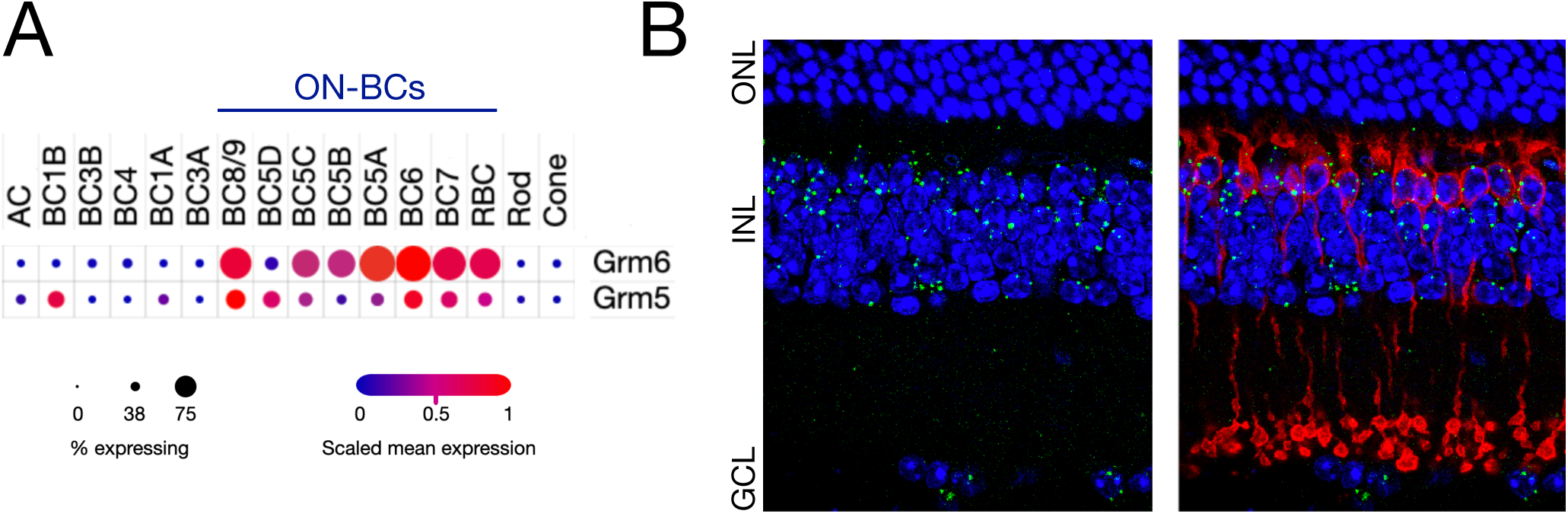
mGluR5 is expressed in bipolar cells. ***A***, mGluR6 and mGluR5 transcripts were detected in retinal bipolar neurons by single-cell RNAseq (adapted from Single Cell Portal, https://singlecell.broadinstitute.org/single_cell, (Tarhan et al., 2023)). ***B***, RNAscope *in situ* hybridization of mouse retinal transversal slices indicate the presence of mGluR5 transcripts in the outer INL (green), where bipolar cell bodies are located, including in rod bipolar cells double-labeled immunohistochemically by PKCα (red). Cell nuclei are stained with DAPI (blue). ONL: outer nuclear layer, INL: inner nuclear layer, GCL: ganglion cell layer. Scale bar: 20 μm.

Significant expression of the *Grm1* gene, encoding mGluR1, was not detected in bipolar cells in these studies (Shekhar et al., 2016; Li et al., 2024). These findings are consistent with several transcriptomic analyses of human retina bipolar cells (Lukowski et al., 2019; Wang et al., 2022), which also show that while mGluR5 is expressed in ON-bipolar cells, mGluR1 expression is restricted to amacrine cells and retinal ganglion cells.

To confirm the presence of mGluR5 transcripts in bipolar cells, we performed RNAscope *in situ* hybridization and co-labelled rod bipolar cells with anti-PKCα antibodies (Fig. 1*B*,*C*). Puncta representing mRNA encoding mGluR5 are notable in the outer half of the inner nuclear layer (INL), where bipolar cell somas are located, with some puncta present in the inner half of the INL, where amacrine cell soma are localized, as well as in the ganglion cell layer (GCL; Fig. 1*B*). Double labeling with anti-PKCα immunohistochemistry to identify rod bipolar cells confirms the presence of mGluR5 in these cells, as well as in cell bodies in the outer INL not labelled by PKCα, presumably cone bipolar cells (Fig. 1*C*).

We then localized Group I receptors in mouse retina using immunofluorescence confocal microscopy. First, we tested several commercial anti-mGluR1 and anti-mGluR5 antibodies on HEK293 cells transfected with cDNAs encoding these receptors. Thus, we identified a monoclonal mouse anti-mGluR1 antibody from BD Bioscience (San Jose, CA; catalog #556389), an anti-mGluR5 antiserum from Spring Biosciences (Pleasanton, CA; catalog #E2870), and a rabbit monoclonal anti-mGluR5 antibody from Cell Signaling (Danvers, MA; catalog #55920) that specifically labeled mGluR1 or mGluR5, respectively. The anti-mGluR5 antiserum was used on an immunoblot with retinal protein extract from wild type and mGluR5^-/-^ mice (Fig. 2*A*). Two bands are visible in the wild type lane, the smaller one corresponding to the receptor monomer, and the larger band corresponding to the dimer, as expected (Romano et al., 1996). Both bands are absent from the mGluR5^-/-^ retinal extracts.

**Figure 2.**
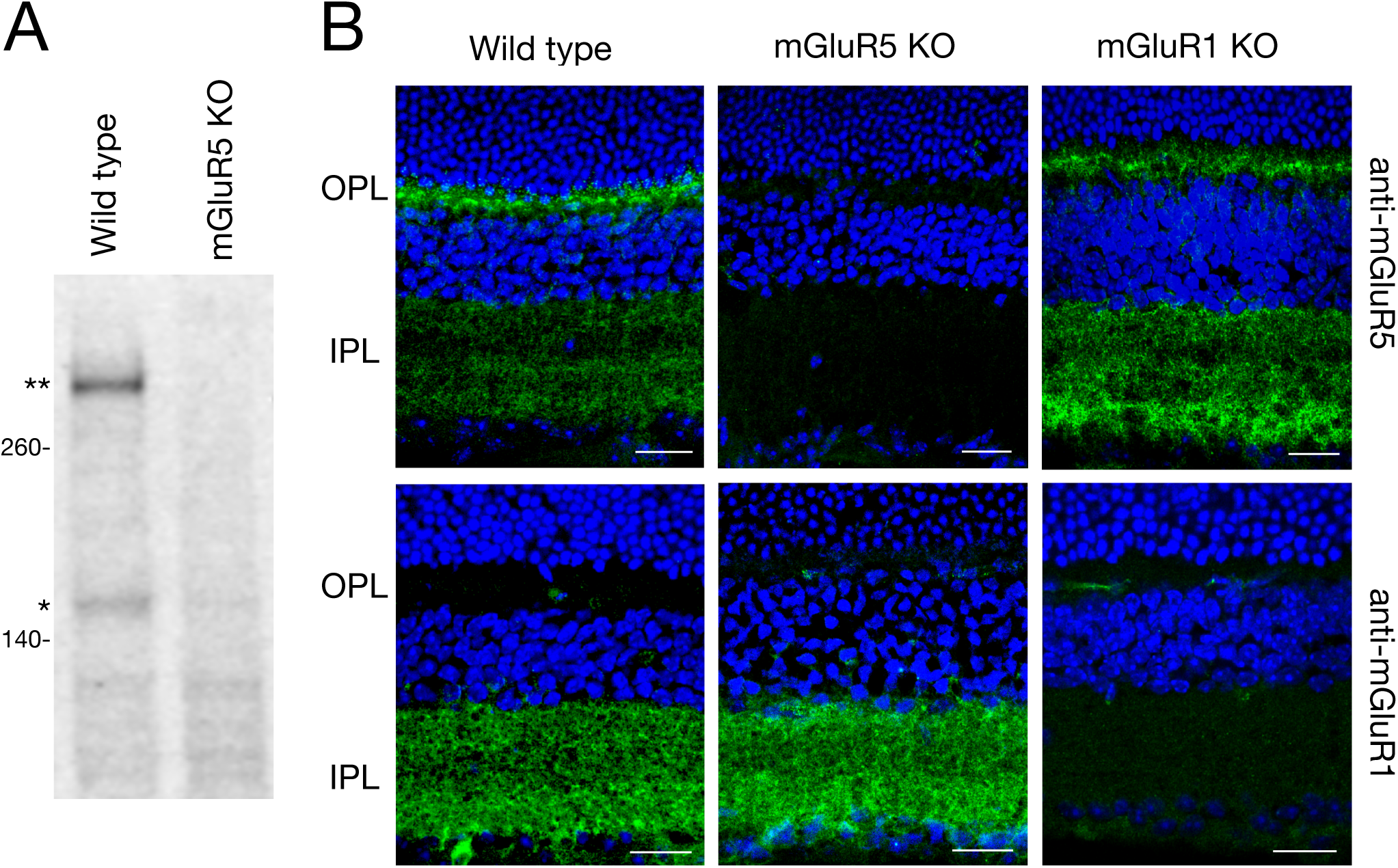
Robust expression of mGluR5, but not mGluR1 in the outer plexiform layer. ***A***, Western blotting reveals monomer (*) and dimer (**) mGluR5 bands in wild type, but not mGluR5 KO retinal protein extracts. ***B***, Immunostaining of retinal sections with anti-mGluR5 antiserum (Spring Biosciences) reveals robust mGluR5 expression in the OPL, and weaker expression in the IPL. Staining is absent in the mGluR5^-/-^ retina. In contrast, mGluR1 expression is confined to the IPL and ganglion cell layer (antibody from BD Biosciences). Cell nuclei are stained with DAPI (blue). OPL: outer plexiform layer, IPL: inner plexiform layer. Scale bars: 20 μm.

These antibodies were validated by immunofluorescent labeling of transverse retina sections from wild type, mGluR5^-/-^, and mGluR1^-/-^ mice (Fig. 2*B*). As shown in the left panels, mGluR5 immunofluorescence is present in both the OPL and IPL. In the OPL (Fig. 2*B* and 3), mGluR5 labeling appears as small puncta, with each punctum presumably corresponding to a pair of rod bipolar cell dendritic tips contacting a rod spherule; as well as larger, densely labelled bars, each presumably corresponding to the numerous bipolar cell dendrites contacting a cone pedicle. Weaker mGluR5 immunofluorescence is present throughout the IPL in the WT retina. In contrast, mGluR1 distribution is restricted to the IPL, presumably on ganglion cell dendrites and amacrine cell processes. Confirming the specificity of the antibodies, no mGluR5 labeling is present in the retina of mGluR5^-/-^ mice and no mGluR1 labeling is visible in mGluR1^-/-^ retinal sections. The remaining fluorescence in mGluR1^-/-^ retina is caused by the rabbit anti-mouse secondary antibody reacting with endogenous mouse IgGs in the vasculature. No changes in mGluR1 expression are detected in mGluR5^-/-^ retina, and reciprocally, the pattern of mGluR5 expression is not altered in the retina of mGluR1^-/-^mice, though the intensity may be increased in the IPL.

**Figure 3.**
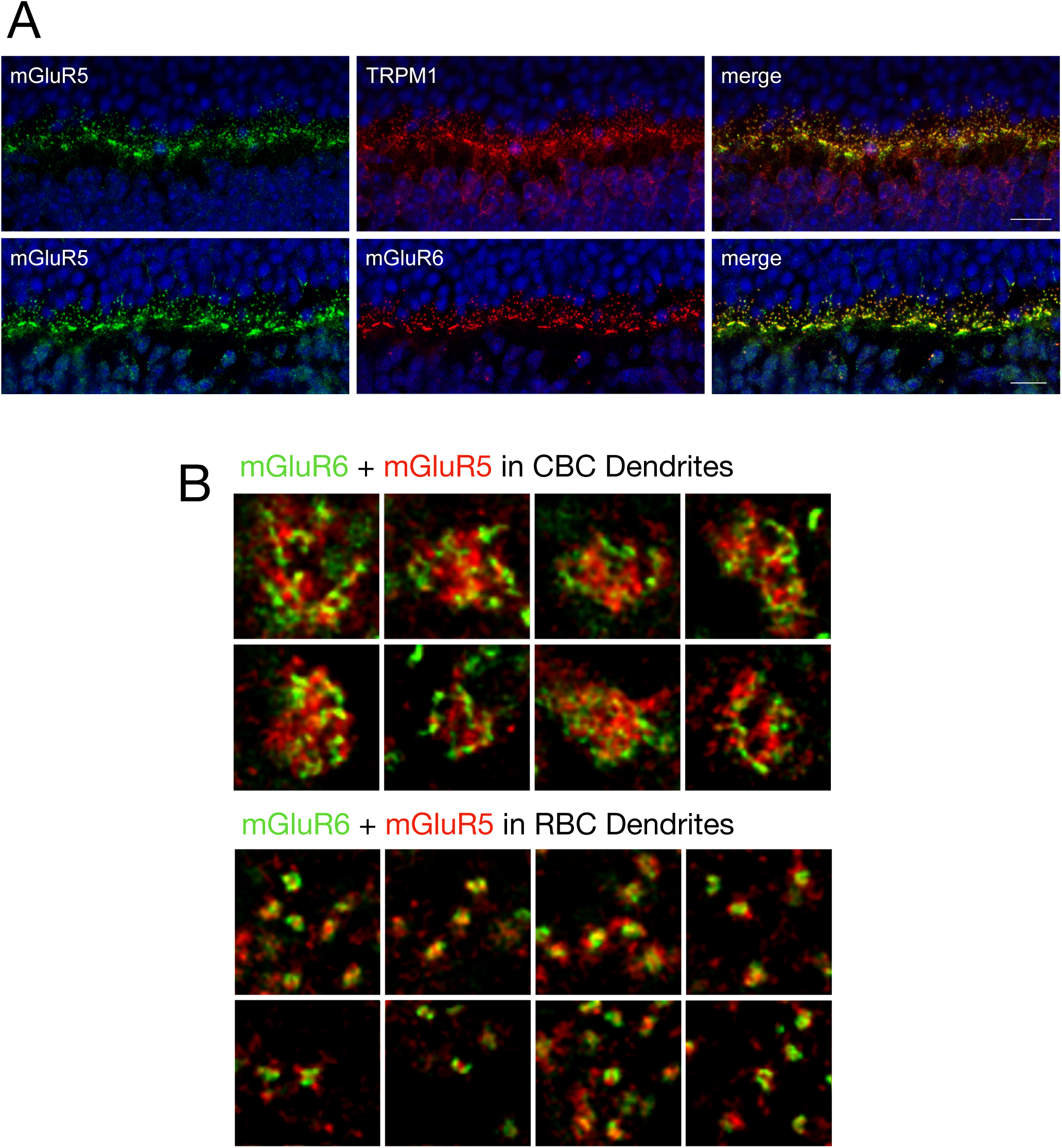
mGluR5 and mGluR6 are located close to one another in ON-bipolar cell dendrites. ***A***, double labeling immunofluorescence of mGluR5 (green, left) and TRPM1 or mGluR6 (red, center), demonstrate mGluR5 localization in the dendritic tips of rod bipolar cells and ON-cone bipolar cells in the merged channels (yellow, right). ***B***, high resolution, confocal images of mGluR5 (green) and mGluR6 (red) double labeling associated with cone and rod synaptic terminals in mouse retina. Scale bars in ***A***: 10 μm. The width of each image in ***B*** corresponds to 5 μm.

Double-labeling immunohistochemistry with antibodies against mGluR5 and either mGluR6 or TRPM1 demonstrates that mGluR5 is localized in the dendrites of rod and cone ON-bipolar cells, together with mGluR6 and TRPM1 (Fig. 3*A*). High-resolution images of mGuR5 and mGluR6 double labeling in the OPL (Fig. 3*B*) reveal that, though the two receptors are located close together, there is little co-localization in either rod or cone ON-bipolar cell dendrites (Pearson’s coefficients: cone bipolar cell dendrites = 0.381 ± 0.151; rod bipolar cell dendrites = 0.514 ± 0.033). mGluR6 is localized to the tips of ON-bipolar cell dendrites that lie within the invaginations of the photoreceptor synaptic terminals (Nomura et al., 1994); thus, mGluR5 is likely to be located in a region of the dendrites more proximal to the ON-bipolar cell bodies.

### The scotopic ERG b-wave is reduced in mGluR5^-/-^ mice

To examine the role of mGluR5 in the light response of rod bipolar cells, we compared scotopic full-field electroretinogram recordings from wild type and mGluR5-deficient mice to a range of light intensities (Fig. 4*A*). The electroretinograms at four light intensities from wild type (10 eyes from 6 mice; black) and mGluR5^-/-^ (8 eyes from 5 mice; red) mice are superposed in Fig. 4*B*. The ERG a-waves, which result from the hyperpolarization of photoreceptors to light, are unchanged by the absence of mGluR5 at dim intensities, but are slightly larger in the mGluR5^-/-^ mice at bright intensities, though the difference compared to wild type is not significant (Fig. 4*C*; maximum a-waves WT: -165 ± 22 μV, n=10, KO: -191 ± 25 μV, n=8, p = 0.43). The b-wave amplitudes in the mGluR5^-/-^ mice are smaller than wildtype for all but the brightest intensities, with the largest differences between the knockout and wild type b-waves occurring in the mesopic range between -3 and -1 log (sc cd·s/m^2^) (Fig. 4*C*). To isolate the effect on the rod bipolar cell response, we calculated the ratio of the amplitude of the b-wave, the rise of which results from the depolarization of rod bipolar cells, to the amplitude of the a-wave, which arises from the hyperpolarization of photoreceptors, for light stimuli above -1 log (sc cd·s/m^2^) that produce consistent a-wave recordings. At these light intensities, the b/a ratio is significantly reduced in mGluR5^-/-^ compared to WT mice (Fig. 4*D*).

**Figure 4.**
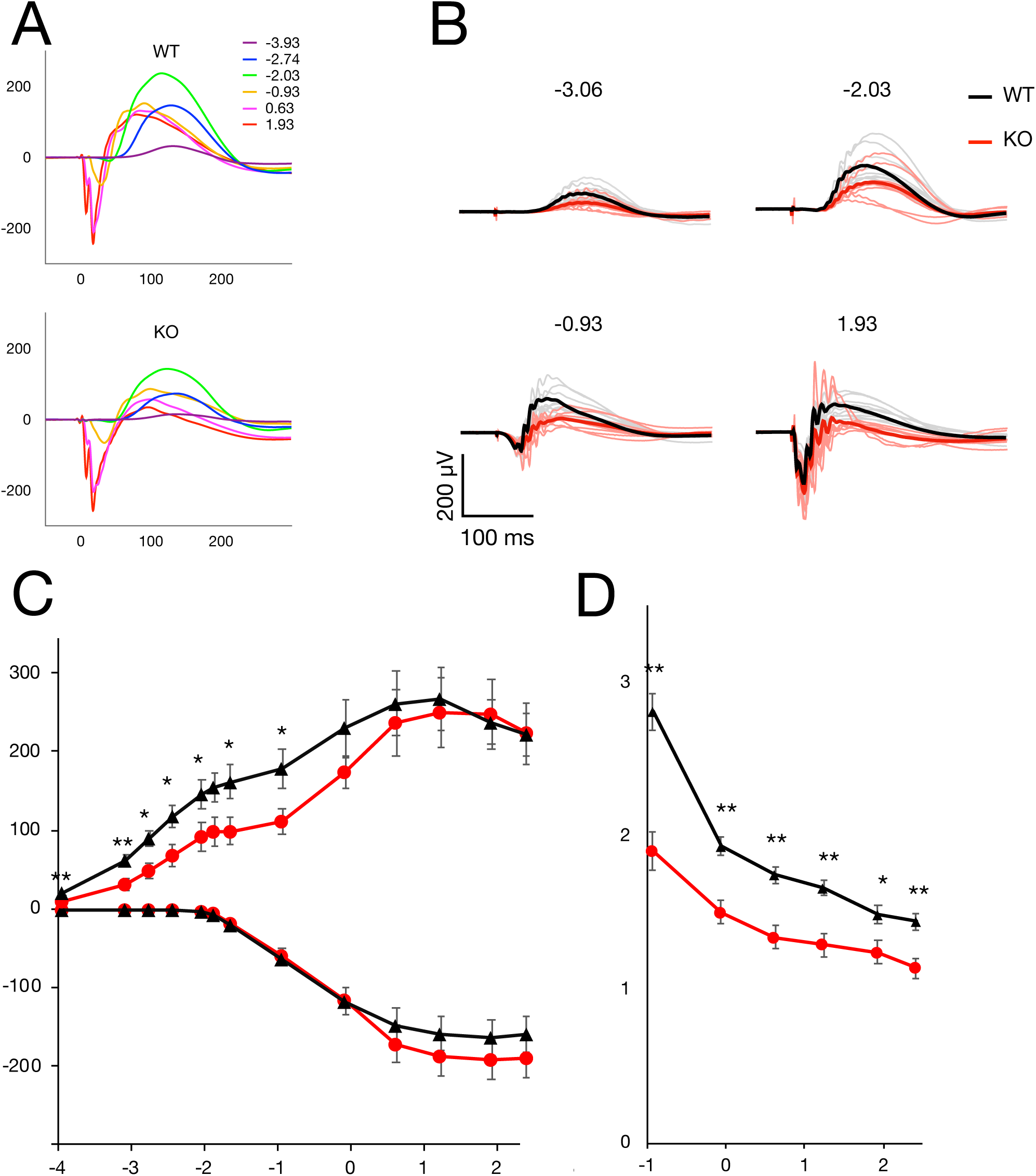
Genetic ablation of mGluR5 reduces the scotopic ERG b-wave amplitude. ***A***, Representative families of scotopic full-field ERG traces to light flashes of increasing intensity from wild type and mGluR5^-/-^ mice. Intensities in log (sc cd•s/m^2^) as indicated by color. ***B***, ERG traces for wild type (black) and mGluR5^-/-^ (red) mice at 4 flash intensities in log (sc cd•s/m^2^). Individual recordings are indicated by fine lines and average of all eyes in bold lines. ***C***, Scotopic a- and b-wave maximal amplitudes of mGluR5 WT (black, n=10) and mGluR5^-/-^ (red, n=8) mice, plotted as a function of flash intensity in log (sc cd•s/m^2^). ***D***, b- to a-wave ratio of mGluR5 WT (black) and KO (red) mice, plotted as a function of flash intensity in log (sc cd•s/m^2^). * p < 0.05, ** p < 0.01.

### Photopic electroretinograms are reduced in mGluR5^-/-^ mice

In addition to expression in rod bipolar cells, single-cell RNAseq studies and our immunohistochemical results show that mGluR5 is present, and possibly even enriched, in cone bipolar cells. To test the function of mGluR5 in cone ON-bipolar cells, we recorded light-adapted, photopic electroretinograms (Fig. 5). At all four stimulation intensities, the amplitudes of the b-wave were lower in mGluR5^-/-^ than in wild type mice (Fig. 5*A*, *B*). Figure 5*C* shows that the average b-wave amplitude is dramatically reduced in the mGluR5^-/-^ compared to WT mice, while the a-wave maximum is only modestly smaller (maximum a-waves WT: -18.4 ± 1.7 μV, KO: -11.6 ± 1.2 μV, p = 0.007; maximum b-waves WT: 82.8 ± 6.0 μV, KO: 33.1 ± 4.3 μV, p = 2·10^-7^). The b/a wave ratio is significantly reduced in mGluR5 KO mice (Fig. 5*D*). Thus, mGluR5 deletion causes an even greater deficit in the function of cone ON-bipolar cells than it does in the function of rod bipolar cells.

**Figure 5.**
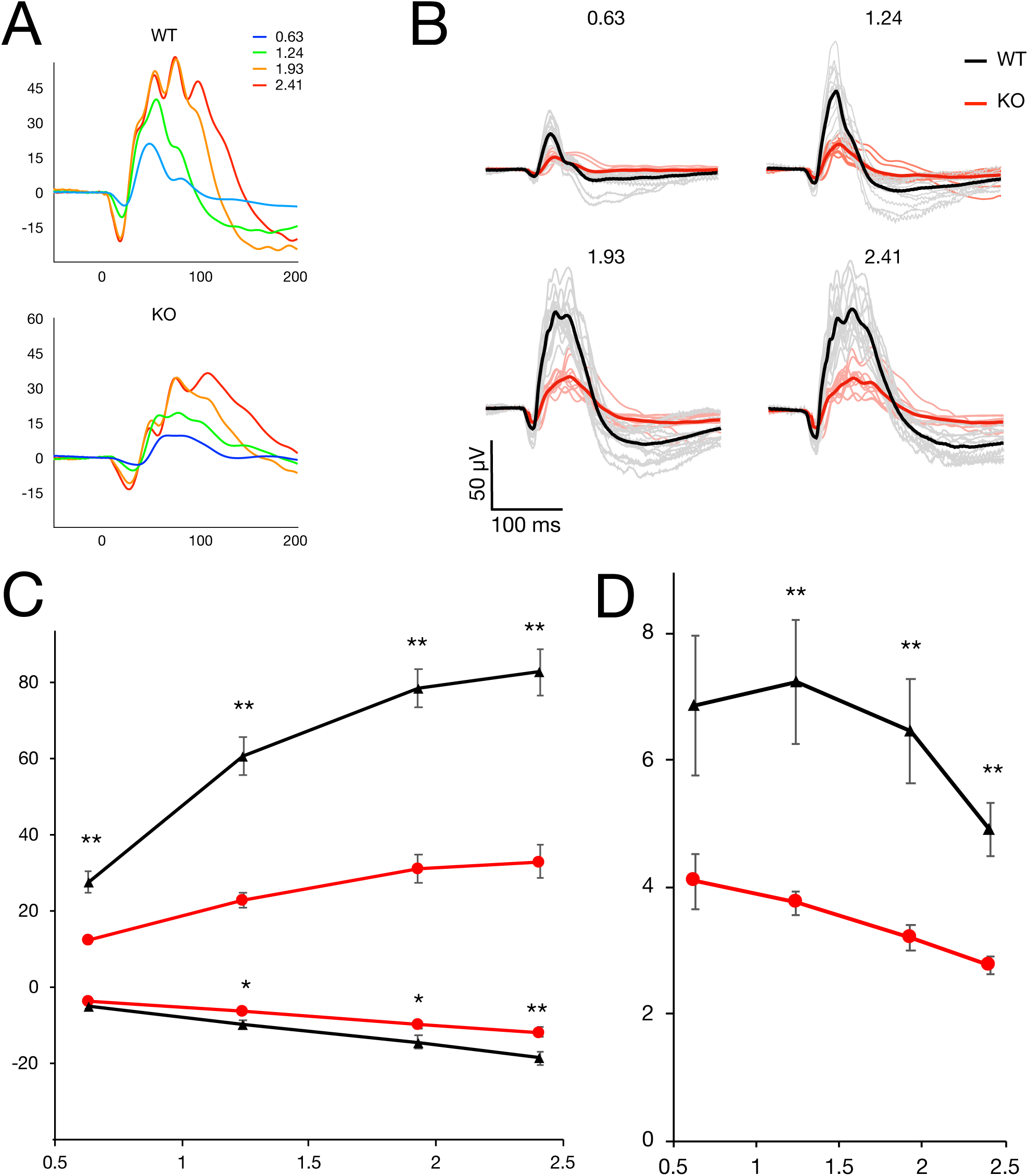
Genetic ablation of mGluR5 reduces the photopic ERG b-wave amplitude. ***A***, Representative families of photopic ERG traces to 4 light flash intensities from wild type and mGluR5^-/-^ mice on a steady 74 cd/m^2^ white light background. Intensities in log (sc cd•s/m^2^) as indicated by color. ***B***, ERG traces for wild type (black) and mGluR5^-/-^(red) mice at 4 flash intensities in log sc cd•s/m^2^. Individual recordings are indicated by fine lines and average of all eyes in bold lines. ***C***, Photopic a- and b-wave amplitudes of mGluR5 WT (black, n=14) and KO (red, n=10) mice, plotted as a function of flash intensity in log (sc cd•s/m^2^). ***D***, b- to a-wave ratio of mGluR5 WT (black) and KO (red) mice, plotted as a function of flash intensity in log (sc cd•s/m^2^). ** p < 0.01.

### Altered localization of ON bipolar cell signaling proteins

To investigate the effect of mGluR5 deletion on other ON-bipolar cell signaling proteins, we examined the distribution of components of the mGluR6 signal transduction pathway by immunofluorescence confocal microscopy (Fig. 6). Transverse retina sections from wild type and knockout mice were prepared in parallel, processed together, and imaged using identical settings. We found that mGluR6 immunolabeling was reduced in the dendritic tips of ON-bipolar cells in mGluR5^-/-^ mice compared to wild type and that soma labeling was increased, suggesting a redistribution of mGluR6 (Fig. 6*A*). The ratio of mGluR6 labeling intensity between the soma (Fig. 6*A*, oINL) and the dendrites (Fig. 6*A*, OPL) was significantly increased in mGluR5^-/-^ mice compared to wild type (WT: 0.22 ± 0.03, n=6; mGluR5^-/-^: 0.45 ± 0.02, n=5; p<0.001). This cell soma labeling was especially apparent in bipolar cells that were not co-labeled with the rod bipolar cell marker PKCα, that is, cone bipolar cells (Fig. 6*B*, asterisks). Immunolabeling for TRPM1, RGS11 and RGS7 was also notably reduced in mGluR5^-/-^ retina (Fig. 6*A*).

**Figure 6.**
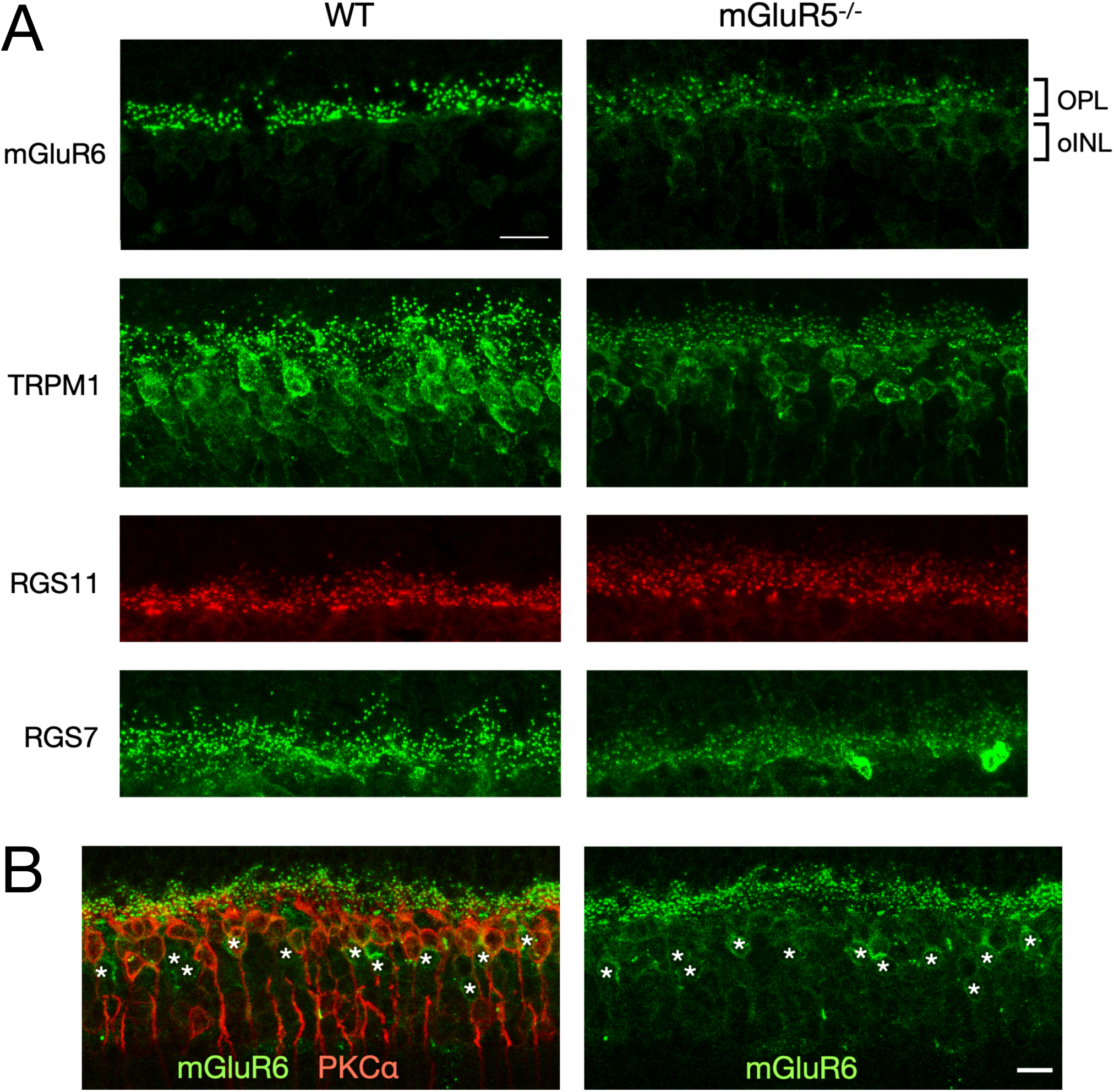
Retinal mGluR6 signaling pathway is affected in mGluR5^-/-^ mice. Transversal sections from wild type (WT) and mGluR5^-/-^ mouse retina were immunolabelled for mGluR6, TRPM1, RGS11 and RGS7. ***A***, Punctate mGluR6 labeling of the ON-bipolar cell dendritic tips in the OPL is reduced, whereas immunolabeling of cell bodies in the outer part of the INL (oINL) is increased in mGluR5^-/-^ retina. TRPM1and RGS7 labeling is reduced in mGluR5^-/-^ retina. RGS11 labeling of ON-cone bipolar cell dendrites appears much reduced in mGluR5^-/-^ retina. OPL: outer plexiform layer; oINL: outer half of inner plexiform layer. ***B***, Redistribution of mGluR6 in the retina of mGluR5^-/-^ mice. Double labeling immunohistochemistry of mGluR6 (green) and PKCα (red) reveal localization of mGluR6 to bipolar cell bodies, especially cone ON-bipolar cells, negative for PKCα, as indicated by *. Scale bars: 20 μm (***A***), 10 μm (***B***).

The ratio of background-corrected total cell fluorescence for identically sized ROIs of mGluR5^-/-^ vs WT was 0.53 ± 0.05 for TRPM1 (n=8, p < 0.001), 0.78 ± 0.08 for GPR179 (n=7, p < 0.05), 0.72 ± 0.05 for RGS11 (n=6, p < 0.01), and 1.01 ± 0.10 (no change) for RGS7 (n=7, p = 0.84). The reduction and re-localization of mGluR6 signal transduction proteins in the ON-bipolar cell dendrites of mGluR5^-/-^ mice (Fig. 6) suggests that mGluR5 is an integral part of the ON-bipolar cell signalosome and contributes to metabotropic glutamate signaling in ON-bipolar cells.

### The ERG b-wave is altered in the presence of the mGluR5 allosteric antagonist MTEP

To assess the physiological role of mGluR5 separately from its effects on the localization of mGluR6 signal transduction components, we used a pharmacological approach, injecting the mGluR5 allosteric antagonist, MTEP (Cosford et al., 2003), to acutely inhibit mGluR5 receptors in adult, wild type mice. First, to establish a time course of the effects of MTEP on the ERG, responses to -0.93 log (sc cd•s/m^2^) light flashes, repeated every 5 min, were recorded following the intra-peritoneal injection of 3 mg/kg MTEP or carrier control. The maximal effect of MTEP was reached by 20 min, compared to the control. Subsequently, ERGs were recorded to the full series of light intensities 20 min after MTEP injection (Fig. 7*A*). ERGs at four light intensities from control (10 eyes from 6 mice; black) and MTEP-administered (11 eyes from 6 mice; green) mice are superposed in Figure 7*B*. MTEP caused a statistically significant increase in b-wave amplitudes for all intensities below -1.85 log (sc cd•s/m^2^) (Fig. 7*C*). In addition, it is notable that the b-wave for a flash of -1.64 log (sc cd•s/m^2^) is smaller than that for a dimmer flash of -1.85 log (sc cd•s/m^2^) in the presence of MTEP. In contrast to our findings comparing WT and mGluR5^-/-^ mice, the scotopic ERG a-wave was slightly larger following MTEP injections between -3.06 and -1.85 log (sc cd•s/m^2^).

**Figure 7.**
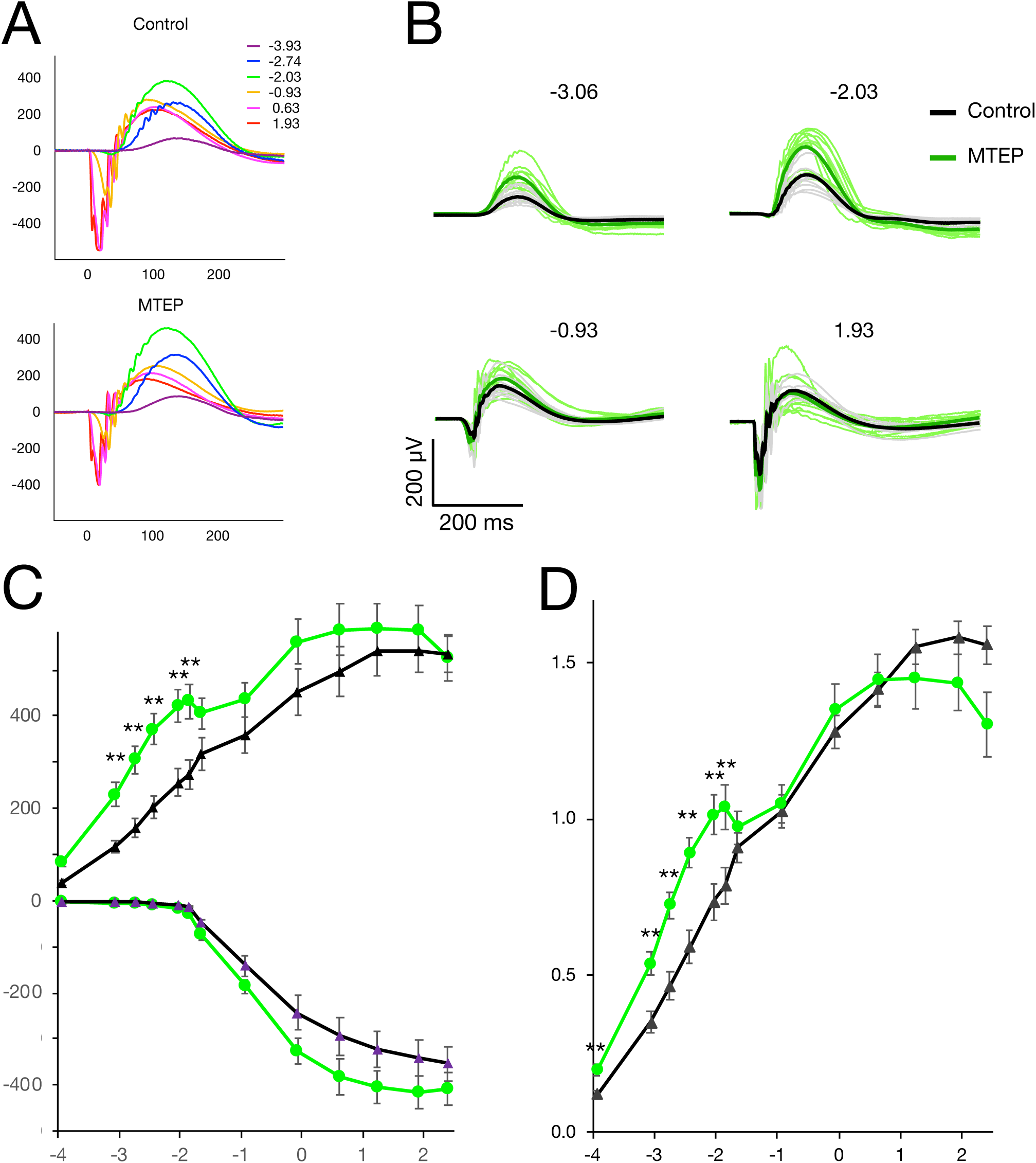
The ERG b-wave is enhanced by the mGluR5 allosteric antagonist MTEP in response to dim flashes. ***A***, Representative families of ERG traces to light flashes of increasing intensity 20 min after s.c. injection of carrier control or 3 mg/kg MTEP. Intensities in log (sc cd•s/m^2^) as indicated by color. ***B***, ERG traces 20 min after s.c. injection of carrier control (black) or 3 mg/kg MTEP (green) at 4 flash intensities in log (sc cd•s/m^2^). Individual recordings are indicated by fine lines and average of all eyes in bold lines. ***C***, Maximal scotopic a- and b-wave amplitudes of control- (black, n=9) and MTEP- (green, n=8) injected mice, plotted as a function of flash intensity in log (sc cd•s/m^2^). ***D***, Normalized b-wave amplitudes of control (black) and MTEP (green), plotted as a function of flash intensity in log (sc cd•s/m^2^). * p < 0.05, ** p < 0.01.

To examine the effect of MTEP on ON-bipolar cells at the dimmest light stimulation intensities when a-waves are too small to be reliably measured, and to remove the contributions of changes in photoreceptor responses, we normalized the b-waves to the largest a-wave of the recording (Fig. 7*D*). Similar to the unnormalized ERGs, MTEP caused a statistically significant increase in normalized b-wave amplitudes up to a flash strength of -1.85 log (sc cd•s/m^2^) and a dip in the intensity-response curve such that the normalized b-wave was larger for a flash strength of -1.85 log (sc cd•s/m^2^) than for the next brighter flash of -1.64 log (sc cd•s/m^2^) (Fig. 7*D*).

### Effects of the CB1R antagonist SR14176A and agonist ACEA on the ERG b-wave

In the brain, it has been shown that postsynaptic mGluR5 receptors mediate a retrograde signal that inhibits glutamate release from the presynaptic axon terminal via dendritic release of the endocannabinoid, 2-AG, which activates presynaptic CB1 receptors (CB1R) (Varma et al., 2001; Ohno-Shosaku et al., 2002; Araque et al., 2017; Augustin and Lovinger, 2018). We speculated that a similar mechanism may occur between ON-bipolar cells and photoreceptors, whereby post-synaptic mGluR5 controls the dendritic release of an endocannabinoid that activates CB1R on the photoreceptor terminals.

Using two monoclonal antibodies to CB1R, both of which have been validated in CB1R-knockout mice (Li et al., 2020; Steiger et al., 2023), we found that CB1R immunofluorescence localizes to both the inner and outer plexiform layers of the retina (Fig. 8*A*), consistent with previous reports (Straiker et al., 1999; Yazulla et al., 1999).

**Figure 8.**
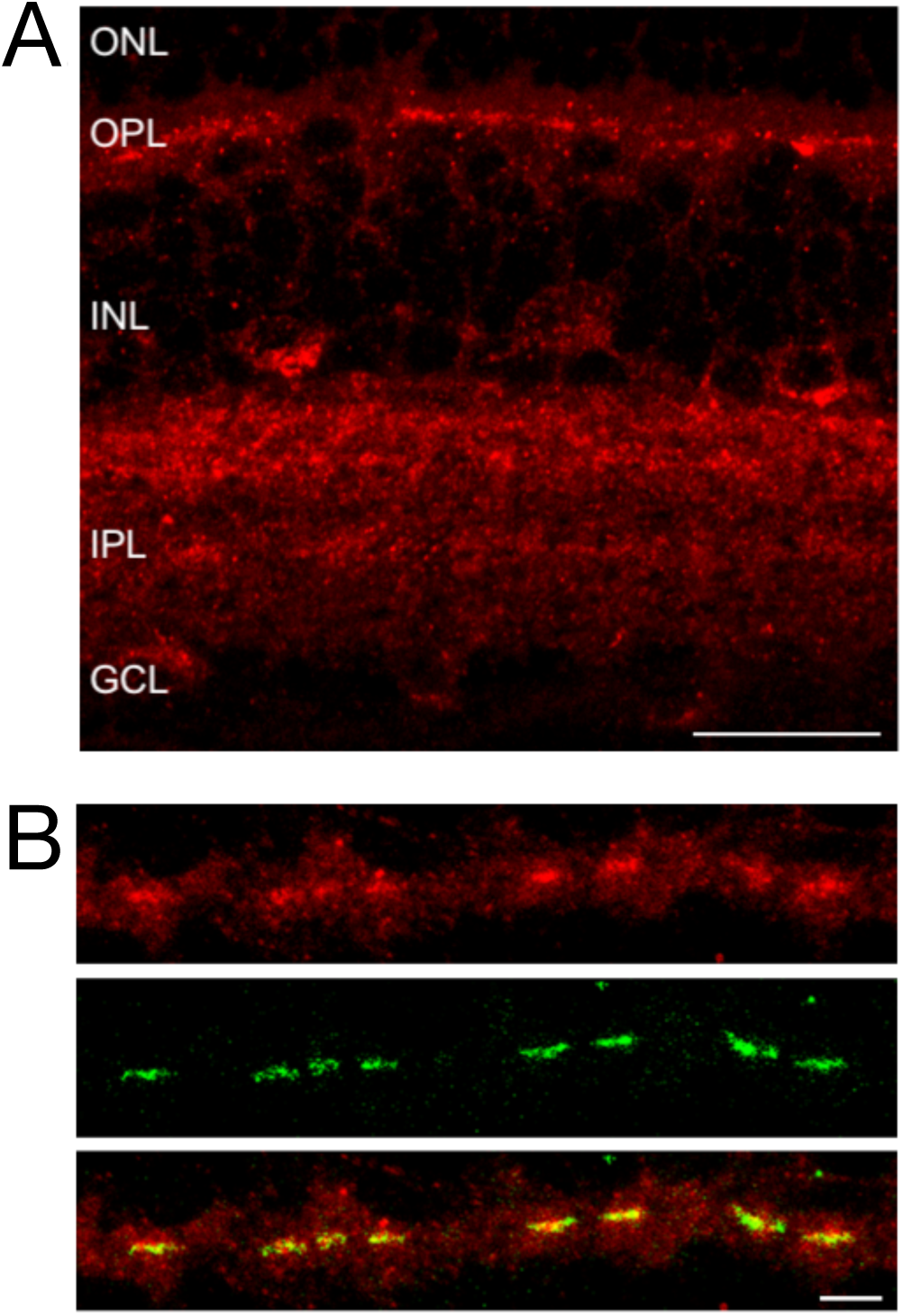
CB1 receptors are expressed in the synaptic layers of the retina. ***A***, Localization of CB1R in the mouse retina (CST antibody). ***B***, Double labeling in the outer plexiform layer for CB1R (red, SySy antibody) and PNA (green), a marker of cone synaptic terminals. The scale bars correspond to 20 μm in ***A*** and 5 μm in ***B***. Abbreviations: ONL, outer nuclear layer; OPL, outer plexiform layer; INL, inner nuclear layer; IPL, inner plexiform layer; GCL, ganglion cell layer.

The two antibodies showed the same distribution in the outer plexiform layer, with the strongest CB1R immunofluorescence coinciding with cone synaptic terminals (Fig. 8*B*), identified by double labeling with PNA, a lectin that binds extracellular sites at cone terminals (Wässle et al., 2009).

To test whether endocannabinoids could affect transmission between photoreceptors and bipolar cells, we recorded ERGs 1 hr following a subcutaneous injection of either the CB1R antagonist/inverse agonist, SR14176A (Fig, 9*A* and *B*), or the CB1R agonist, ACEA (Fig. 9*C* and *D*). Similar to the effect of MTEP, SR14176A caused a significant increase in b-wave amplitudes for all intensities below -1.85 log (sc cd•s/m^2^) (Fig. 9*A*; control: 14 eyes from 7 mice, black; SR14176A: 13 eyes from 7 mice, blue). Notably, the b-waves for flashes of -1.64 and -0.93 log (sc cd•s/m^2^) are smaller than those for dimmer flashes of -1.85 and -2.03 log (sc cd•s/m^2^). Normalizing the ERG b-waves to the largest a-wave of each recording (Fig. 9*B*) confirmed the increased b-wave at dim light intensities, ≤ -1.85 log (sc cd•s/m^2^). In the mesopic range, between -1.64 to -0.93 log (sc cd·s/m^2^), the normalized b-waves of SR14167A injected mice were smaller than at dimmer intensities, similar to the effect of MTEP in this intensity range (Fig. 7*D*). In addition, the normalized b-wave amplitude was reduced at the brightest light stimulations (≥ 1.93 log (sc cd·s/m^2^)) in SR14167A-injected mice compared to controls (Fig. 9*B*), a trend also seen in the MTEP-injected mice (Fig. 7*D*).

**Figure 9.**
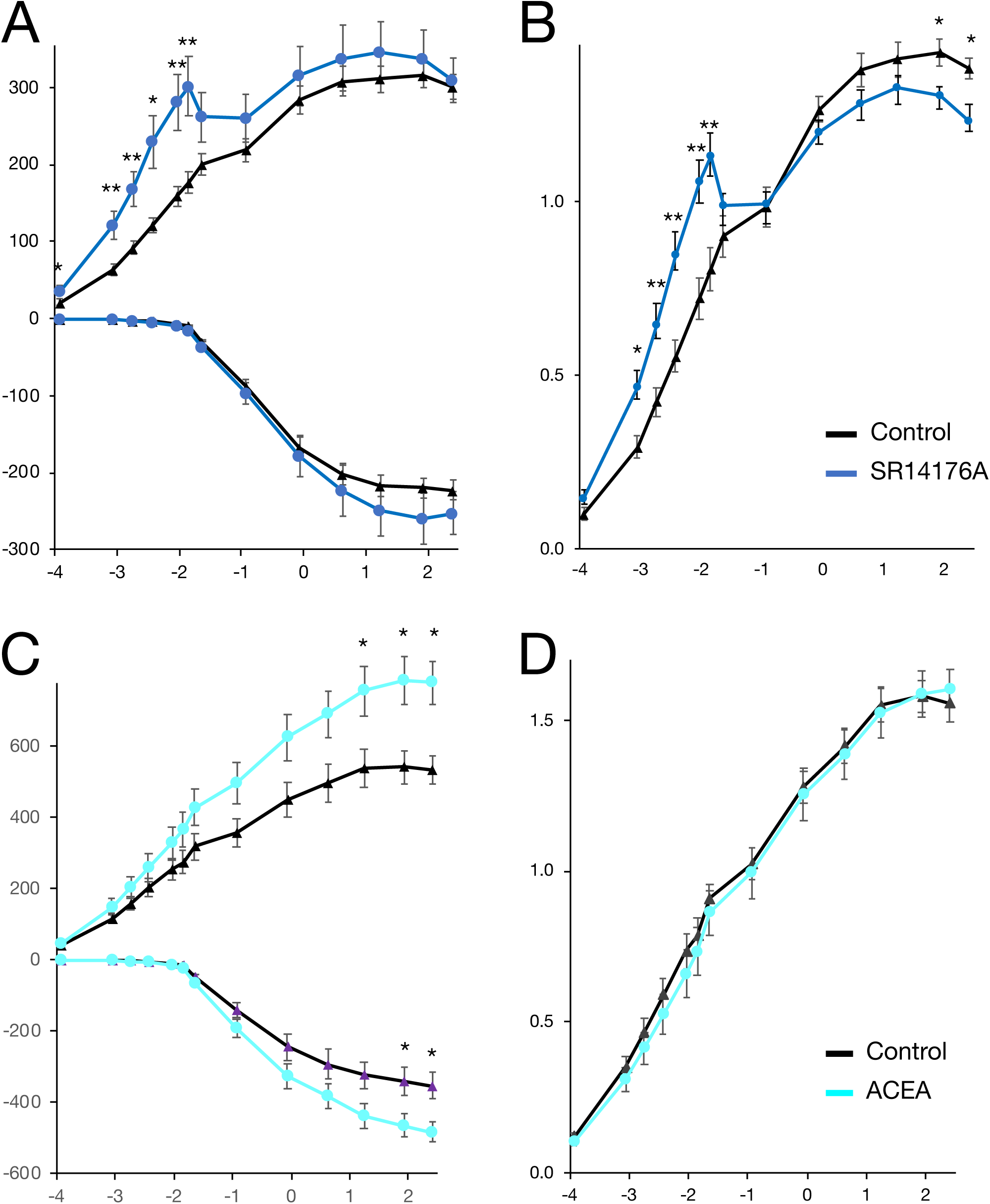
Effects of the CB1 allosteric antagonist SR14176A and the agonist ACEA on the ERG b-wave. ***A***, Scotopic a- and b-wave amplitudes 60 min after s.c. injection of control (black, n=14) or 3 mg/kg SR14176A (blue, n=13), plotted as a function of flash intensity in log (sc cd•s/m^2^). ***B***, Normalized b-wave amplitudes of control (black) and SR14176A (blue), plotted as a function of flash intensity in log (sc cd•s/m^2^). ***C***, Scotopic a- and b-wave amplitudes 30 min after s.c. injection of control (black, n=9) or 1 mg/kg ACEA (cyan, n=10), plotted as a function of flash intensity in log (sc cd•s/m^2^). ***D***, Normalized b-wave amplitudes of control (black) and ACEA (cyan), plotted as a function of flash intensity in log (sc cd•s/m^2^). * p < 0.05, ** p < 0.01.

The CB1 agonist ACEA did not affect ERG responses to dim stimuli; however, both the a- and b-waves were significantly increased at bright stimuli (above 1.93 log (sc cd·s/m^2^) for a-waves and above 1.24 log (sc cd·s/m^2^) for b-waves) as shown in Figure 9*C* (control: 9 eyes from 5 mice, black; ACEA: 10 eyes from 5 mice, cyan). Upon normalizing the ERG b-waves to the largest a-wave of each recording (Fig. 9*D*), there is no effect of ACEA, and it is apparent that the larger non-normalized b-waves in the presence of ACEA (Fig. 9*C*) are due to enlarged inputs from the photoreceptors, as suggested by the larger a-waves (Fig 9*D*).

### The CB1 agonist ACEA reverses the effect of the mGluR5 antagonist MTEP at dim light intensities

To test if CB1R is downstream of a mGluR5-driven retrograde signal from ON bipolar cells to photoreceptors, we compared ERGs recorded following administration of MTEP alone, ACEA alone, MTEP plus ACEA, or the carrier control (Fig. 10). In contrast to MTEP alone, the b-wave of MTEP+ACEA is not significantly different from that of control (Fig. 10*A*). This effect is more noticeable when the ERG b-waves are normalized to the largest a-waves (Fig. 10*B*). Thus, the increased b-wave at dim light intensities and the inverted stimulus-response relationship at mesopic intensities produced by antagonizing mGluR5 is reversed by the CB1 agonist.

**Figure 10.**
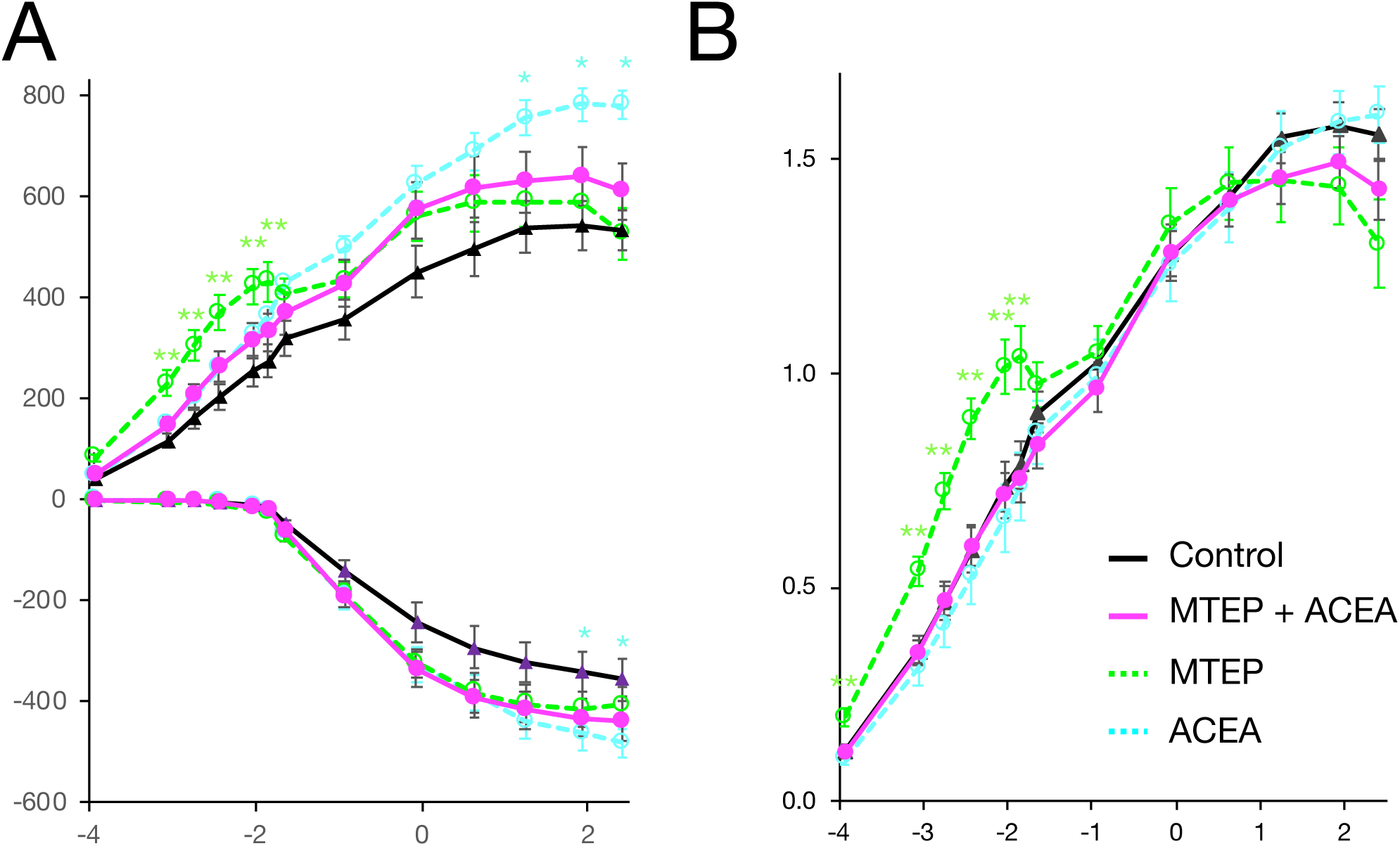
The CB1 agonist ACEA reverses the effect of the mGluR5 antagonist MTEP at dim light intensities. ***A***, Maximal scotopic a- and b-wave amplitudes of control- (black, n=9) and MTEP + ACEA (purple, n=14) -injected mice, plotted as a function of flash intensity in log (sc cd•s/m^2^). For comparison, the a- and b-wave amplitudes for each drug alone, shown in Figures 7C and 9*C* are indicated in dotted lines (MTEP, green; ACEA, cyan). ***B***, Normalized b-wave amplitudes of control (black) and MTEP + ACEA (purple), plotted as a function of flash intensity in log (sc cd•s/m^2^). For comparison, the normalized b-wave amplitudes for each drug alone, shown in Figures 7D and 9*D* are indicated in dotted lines (MTEP, green; ACEA, cyan).

## Discussion

Here we show that mGluR5 receptors are present in the dendritic tips of rod and cone ON-bipolar cells, where they regulate ON light responses, as measured by ERG recording. Several groups have localized either mGluR1, mGluR5, or both, in ON-bipolar cell dendrites, in addition to mGluR6, using immunohistochemical or electrophysiological methods (Koulen et al., 1997; Rampino and Nawy, 2011; Hellmer et al., 2018). Recent single-cell RNA-sequencing analyses from human and mouse retina indicate that mGluR5 is expressed in bipolar cells, whereas mGluR1 is expressed in amacrine and ganglion cells (Fig. 1A; Shekhar et al., 2016; Li et al., 2024; Lukowski et al., 2019; Wang et al., 2022). Using RNAscope, an amplified in situ hybridization method, combined with immunohistochemistry, we confirm that mGluR5 transcripts are found in rod bipolar cells as well as putative cone bipolar cells and ganglion cells (Fig. 1*B*).

Using antibodies specific for either mGluR1 or mGluR5, we demonstrate that mGluR5 is found in both the outer and inner plexiform layers, but mGluR1 is only detected in the IPL, and not in the OPL. This finding is inconsistent with several previous studies that have shown mGluR1 labeling in the OPL of rat and mouse retinas (Koulen et al., 1997; Hellmer et al., 2018), but our results are in agreement with Tagawa et al. (1999) and Hozumi et al. (2013), who also immunolocalized mGluR5 to the OPL and IPL, but mGluR1 only to the IPL of mouse and rat retinas. In seeking antibodies specific for mGluR5 and mGluR1, we tested antibodies on transfected HEK293 cells and found that many commercial antibodies cross-reacted with both receptors, possibly explaining erroneous detection of mGluR1 in the OPL in some of the earlier studies.

Double labeling experiments localized mGluR5 receptors to the dendrites of rod and cone ON-bipolar cells, in close proximity to mGluR6 and TRPM1 (Fig. 3). It is known that mGluRs can form heterodimers. This has been shown for the group I, mGluR1 and mGluR5 receptors, as well as within and between group II and III mGluRs (Doumazane et al., 2011). However, formation of heterodimers between group I and the other groups has not been reported. As there is little overlap between the mGluR5 and mGluR6 immunolabeling patterns (Fig. 3 *B*), it is unlikely that they form heterodimers in ON-bipolar cells. Furthermore, the ERGs of mGluR5^-/-^ mice indicate that heterodimerization of mGluR6 and mGluR5, if present, is not required for the ON-bipolar cell light response, as the mGluR5 knockout mice generate a robust, albeit reduced, b-wave (Fig. 4). Instead, their close, but non-overlapping proximity in ON-bipolar cell dendrites suggests they respond independently to glutamate release from photoreceptors. The relative distributions of the two receptors are consistent with the localization of mGluR6 to the ON-bipolar cell dendritic tips and of mGluR5 in the dendrites, but more proximal to the ON-bipolar cell bodies, with mGluR5 likely residing outside the synaptic invaginations of the photoreceptor terminals, similar to its perisynaptic localization in the brain (Lujan et al., 1996). Such an arrangement implies that mGluR5 would likely experience lower glutamate concentrations than mGluR6, since photoreceptors release glutamate into the extracellular space within the invaginations.

If mGluR5 signaling in ON-bipolar cells utilizes a similar downstream pathway as mGluR6, one might expect an mGluR5-driven b-wave to be present in the absence of mGluR6, but this is not seen in mGluR6^-/-^ mice. Genetic deletion of mGluR6, however, results in the loss of other components of the signaling complex from the ON-bipolar cell dendrites, including TRPM1 (Morgans et al., 2007; Xu et al., 2012), and also eliminates the trans-synaptic connection with the presynaptic membrane that is mediated by binding between mGluR6 and ELFN adhesion proteins (Cao et al., 2015, 2020; Miller et al., 2024). Thus, knockout of mGluR6 has consequences beyond the elimination of mGluR6-driven signaling that would be expected to also impair mGluR5 signaling.

Similarly, we show that knockout of mGluR5 is associated with a decrease in and/or mislocalization of key proteins at the tips of ON-bipolar cells dendrites, including mGluR6 and TRPM1 (Fig. 6). Notably, mGluR5 has been shown to interact with Homer and other postsynaptic density proteins (Xiao et al., 2000), with several of these proteins expressed in ON-bipolar cell dendrites (Brandstätter et al., 2004). Thus, the reduction in the amplitude of the ERG b-wave across all light intensities in the mGluR5^-/-^mice might be explained in part by the effect of the mGluR5 knockout on the expression of other signaling proteins.

The relative effects of mGluR5 and mGluR6 on ON-bipolar responses to changing luminance are likely to be influenced by their binding affinities for glutamate. The EC_50_ for glutamate is reported to be 3–10 μM for rat mGluR5 and 15–40 μM for rat mGluR6 (Schoepp et al., 1999). Therefore, it is possible that small decreases in synaptic glutamate, such as those resulting from single photon absorption by rods in a dark-adapted retina, would be transmitted to rod bipolar cells via mGluR6, but that synaptic glutamate levels would not decrease sufficiently to cease to stimulate mGluR5. Brighter light flashes would be expected to decrease activity of both mGluR6 and mGluR5.

The signaling pathways and time courses of mGluR6 and mGluR5 may also differ. In response to light increments, ON-bipolar cells depolarize within tens of milliseconds (Berntson and Taylor, 2000; Sampath and Rieke, 2004) as TRPM1 channels open when their inhibition by Go is relieved, following the hydrolysis of Gαo-GTP and the reformation of Gαo-GDP-βγ trimers. Recent experiments based on bioluminescent resonance energy transfer (BRET) suggest that mGluR5 can also activate the Go pathway (Masuho et al., 2023), but mGluR5 is generally thought to be coupled to Gq and phosphoinositide signaling pathway molecules (Conn and Pin, 1997), such as phospholipase C (PLC) β, diacylglycerol kinase (DGK) and PKCα, which have all been shown to be localized in the OPL (Hozumi et al., 2013). These pathways are slower and, in the retina, may be able to integrate light intensity over a few minutes (Rampino and Nawy, 2011; Hellmer et al., 2018).

Similar to many GPCRs, mGluR5 rapidly desensitizes upon prolonged exposure to agonist partially as a result of phosphorylation by PKC or a GPCR kinase (Schoepp and Johnson, 1988; Catania et al., 1991; Gereau and Heinemann, 1998; Sorensen and Conn, 2003; Gainetdinov et al., 2004; Lee et al., 2008; Ko et al., 2012), whereas mGluR6 does not desensitize. Thus, in the dark, mGluR6 stimulation is expected to be continuous and TRPM1 channels remain closed; whereas mGluR5 might be desensitized and no longer coupled to G proteins. Upon light exposure and decreased glutamate release from photoreceptors, mGluR5 receptors are expected to recover from desensitization within 30 min (Gereau and Heinemann, 1998), so that a return to darkness would stimulate the receptors. In this way mGluR5 responses would be transient, depending on the history of light exposure and may serve to preserve a “memory” of the recent history of light exposure. Such a mechanism may contribute to light adaptation of photoreceptor to bipolar cell transmission.

In the brain, post-synaptic mGluR5 has been linked to retrograde endocannabinoid signaling through its role in stimulating synthesis of the endocannabinoid, 2-AG (Varma et al., 2001; Ohno-Shosaku et al., 2002; Augustin and Lovinger, 2018). 2-AG is an important mediator of synaptic suppression at many brain circuits, generally acting via presynaptic CB1 and CB2 cannabinoid receptors on voltage-gated calcium channels and G protein inwardly rectifying potassium channels to reduce neurotransmitter release (Araque et al., 2017). Gq-coupled, group I mGluRs activate phospholipase C, which hydrolyses phosphatidylinositol 4,5-biphosphate (PIP_2_) into inositol triphosphate (IP_3_) and diacylglycerol (DAG). In turn, membrane-bound diacylglycerol lipase hydrolyses arachidonic acid-containing DAG to 2-AG and a fatty acid. Single-cell RNA-seq experiments have shown the expression of diacylglycerol lipase A and B in bipolar cells and CB1 receptors in photoreceptors (Shekhar et al., 2016; Li et al., 2024). In addition, we and others have immunolocalized CB1 receptors to the OPL (Fig. 8: Straiker et al., 1999; Yazulla et al., 1999), most prominently in cone terminals (Fig. 8*B*).

Thus, we considered whether mGluR5 in ON-bipolar cells could modulate photoreceptors to bipolar cell transmission via dendritic release of 2-AG, which would diffuse trans-synaptically to stimulate CB1 receptors on photoreceptor terminals. We tested the effect of the CB1 antagonist, SR14176A, on the mouse full-field ERG and found that it caused a substantial increase in the amplitude of the ERG b-wave for dim flashes (≤ -1.85 log (sc cd·s/m^2^), Fig. 9*A*,*B*), similar to the effects of the mGluR5 antagonist, MTEP (Fig. *7C*,*D*), whereas the CB1 agonist, ACEA, had no significant effect (Fig. 9*D*). These results suggest that mGluR5 may modulate glutamate release for dim flashes via retrograde stimulation of photoreceptor CB1 receptors. As the flash intensity increased, the CB1 antagonist, SR14176A, caused a sudden drop in the b-wave amplitude for flashes in the mesopic range (between -2 and 0 log (sc cd·s/m^2^) (Fig. 9*A,B*), again similar to that observed with MTEP (Fig. 7*D*). This dramatic drop in the response-intensity relationship around -2 log (sc cd·s/m^2^) is also similar to the reduction in the mesopic b-wave amplitude observed in connexin 36 knockout mice, which lack gap junctions between rod and cone terminals and, thus, the secondary rod pathway (Abd-El-Barr et al., 2009; Pasquale et al., 2020). In goldfish, CB1 receptors have been shown to modulate electrical coupling between rods and cones (Cao and Mangel, 2021). Thus, at mesopic light intensities that drive mGluR5 signaling, 2-AG released from ON-bipolar cells could function as a retrograde messenger to modulate photoreceptor electrical coupling.

To further test whether stimulation of mGluR5 on ON-bipolar cell dendrites leads to a cannabinoid-driven retrograde signal onto photoreceptors, we recorded the ERGs of mice administered both the mGluR5 antagonist MTEP and the CB1 receptor agonist ACEA (Fig. 10). We found that ACEA reversed the effects of MTEP on the b-wave amplitudes at dim light stimulations, as well as the dip in the response-intensity relationship observed under mesopic light stimulations, whereas ACEA alone had no effect on the normalized b-wave under these light stimulation conditions. Thus, CB1 receptors are physiologically located downstream of mGluR5 and mediate its regulation of synaptic transmission between photoreceptors and ON-bipolar cells in the retina.

Taken together, our data demonstrate a role for mGluR5 in regulating synaptic transmission from photoreceptors to ON-bipolar cells. Possible downstream pathways include modulation of TRPM1 and retrograde, trans-synaptic endocannabinoid signaling that could regulate glutamate release from photoreceptors as well as gap junction coupling between rods and cones.

## Acknowledgments

We thank Elisa Guimaraes-Souza for help finding the mGluR1 and mGluR5 specific antibodies, Louis Duvoisin for help writing Python scripts, Allura Thompson and Hope Shi for help with genotyping and immunohistochemistry quantifications, respectively. We thank Geoff Swanson (Northwestern), Kimberly Huber (UT Southwestern) and François Conquet (Exciva GmbH, Heidelberg) for providing the mGluR1 and mGluR5 KO mice. We thank Lane Brown (Washington State University) for advice and discussions.

Supported in part by NEI grant R01 EY029985

